# Global analysis of protein stability by temperature and chemical denaturation

**DOI:** 10.1101/2020.04.20.049429

**Authors:** Louise Hamborg, Emma Wenzel Horsted, Kristoffer Enøe Johansson, Martin Willemoës, Kresten Lindorff-Larsen, Kaare Teilum

## Abstract

The stability of a protein is a fundamental property that determines under which conditions, the protein is functional. Equilibrium unfolding with denaturants requires preparation of several samples and only provides the free energy of folding when performed at a single temperature. The typical sample requirement is around 0.5 – 1 mg of protein. If the stability of many proteins or protein variants needs to be determined, substantial protein production may be needed. Here we have determined the stability of acyl-coenzyme A binding protein at pH 5.3 and chymotrypsin inhibitor 2 at pH 3 and pH 6.25 by combined temperature and denaturant unfolding. We used a setup where tryptophan fluorescence is measured in quartz capillaries where only 10 μl is needed. Temperature unfolding of a series of 15 samples at increasing denaturant concentrations provided accurate and precise thermodynamic parameters. We find that the number of samples may be further reduced and less than 10 μg of protein in total are needed for reliable stability measurements. For assessment of stability of protein purified in small scale e.g. in micro plate format, our method will be highly applicable. The routine for fitting the experimental data is made available as a python notebook.

## Introduction

The stability of the native state of a protein determines under which conditions the protein is folded and thus active. Accurate measurements of protein stability are important if we wish to understand the underlying interactions that stabilizes a protein structure and manipulate proteins to be more (or less) stable. Protein stabilities are typically determined by gradually changing temperature or the concentration of a chemical denaturant and measuring the unfolding by a spectroscopic technique or calorimetry. For highly stable proteins, e.g. from thermophilic organisms, proteins may not necessarily unfold at temperatures below 100 °C and it may be thus be necessary to include a chemical denaturant to reach the unfolded state. Similarly, some proteins need elevated temperatures to completely unfold in a chemical denaturant.

Protein denaturation may be measured by differential scanning calorimetry (DSC), where the excess heat capacity of unfolding is quantified, and gives a direct measure of the enthalpy for folding and the melting temperature (Tm) [1]. DSC thus provides a full thermodynamic description of the folding process as long as it is reversible, though fitting DSC experiments and determining the thermodynamic stability at ambient temperatures is not always trivial. Protein folding can also be followed by different types of spectroscopies. Most widely applied are fluorescence spectroscopy and CD spectroscopy [2]. In fluorescence spectroscopy, the process may either be probed by the change in the intrinsic protein fluorescence from tryptophan or tyrosine residues or by the change in fluorescence from extrinsic fluorescent probes such as SYPRO orange and ANS that binds to hydrophobic parts of the protein [3]. The fluorescence signals of fluorophores are highly dependent on the surrounding environment, which can be used as a measure for protein unfolding [4]. The Prometheus nanoDSF apparatus (NanoTemper Technologies GmbH) makes it possible to measure unfolding using small volumes (10 μl) and low concentrations of proteins (down to 5 μg/mL) and allows for the analysis of up to 48 samples simultaneously. The instrument uses quartz capillaries as sample cuvettes and measures the intrinsic protein fluorescence at 330 nm and 350 nm (with excitation at 280 nm) as a function of temperature from 15 °C to 95 °C. It is thus possible to perform unfolding experiments by temperature and denaturant on small amounts of protein.

Here we have performed two-dimensional unfolding experiments on bovine acyl-coenzyme A binding protein (ACBP) and barley chymotrypsin inhibitor 2 (CI2). The folding mechanisms of these proteins have been intensively studied [5,6,15–18,7–14] and ACBP and CI2 thus serve as an excellent set of model proteins for validating our method. By fitting temperature denaturation profiles at several denaturant concentrations to a multivariate function with both temperature and denaturant concentration as independent variables it is possible to estimate Δ*H_m_*, Δ*C*_p_, *T*_m_ and the *m*-values as demonstrated by others [19]. We show that the total number of different GuHCl concentrations needed for a robust parameter estimate can be drastically reduced, and that both *T*_m_ and Δ*H* at 298K still get reliably determined from only five samples for both ACBP and CI2. With the need of only a low amount of protein for each sample and the few samples, the method is well suited if the available amount of protein is limited, and for measuring many variants in parallel.

### Theory

#### Temperature denaturation

In the following, we will consider the denaturation of a protein that (un-)folds reversibly by a two-state process 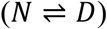. The equilibrium constant *K* for unfolding is then: *K* = [*D*]/[*N*]. *K* will depend on the temperature *T* according to:

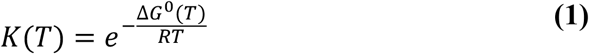

where Δ*G*°(*T*) is the temperature dependent change in Gibbs free energy for unfolding and *R* is the gas constant. The temperature midpoint of the unfolding, *T*_m_, defines the temperature at which the fractions of native and denatured protein are equal (Δ*G*°(*T*_m_) = 0). Δ*G*°(*T*) is given by:

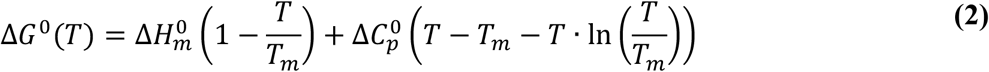

where Δ*H*_m_ is the change in enthalpy upon unfolding at *T*_m_ and 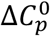 is the change in heat capacity of the system which here is assumed to be independent of temperature [20].

For a two-state process, the fluorescence signal (or any other spectroscopic signal) of a thermally induced denaturation can be described by the equilibrium constant and the temperature dependent fluorescence signals of the native and denatured protein, *f*_N_(*T*) and *f*_D_(T), respectively. Also, for a two-state unfolding process we have that the populations of the two states add to 1, *p*_N_+*p*_D_ = 1. Combining this and using equation 1 we get:

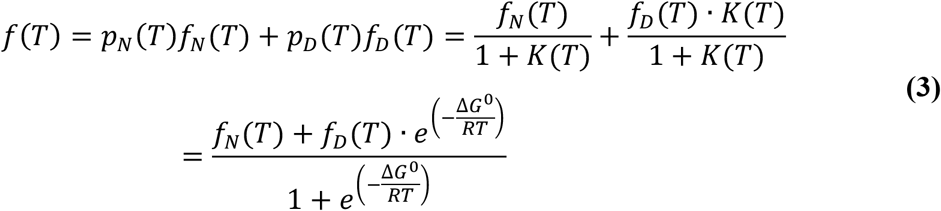

The temperature dependence of the fluorescence signals of the native state is often linear whereas that of the denatured state is curved [21] and often may be described by adding a quadratic term [22], though other non-linear functions have also been used [23]. Thus we have:

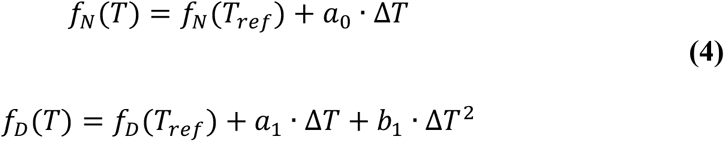

where, *a*_0_, *a*_1_ and *b*_1_ are constants describing the temperature dependence of the fluorescence signal. Δ*T* is defined as Δ*T* = *T* – *T_ref_*, where *T_ref_* is an arbitrary chosen reference temperature for the fluorescence signal.

Introducing the temperature dependence of the fluorescence signals and expanding *DG* according to equation 2 gives:

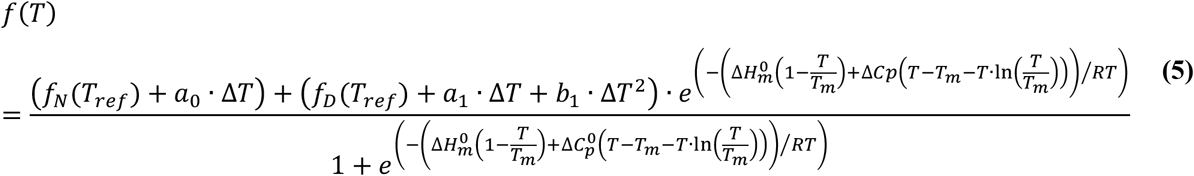

#### Chemical denaturation

Proteins can also be unfolded by denaturants such as urea and guanidinium salts. A number of different models have been described to extract the free energy of unfolding in the absence of denaturant. The most common method is the linear extrapolation model (LEM) [24–26], where the free energy is assumed to be linearly dependent on the denaturant concentration, [x]:

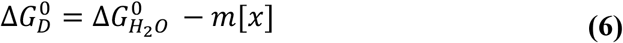

where 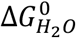 is the free energy of unfolding in the absence of denaturant and *m* describes the (linear) effect of the denaturant on protein stability. The exact mechanistic origin of the *m*-value and equation 6 is not clear, and other methods have tried to rationalize the size of the denaturant dependence. These include the denaturant binding model which assumes that the denaturant binds to sites on the protein and that the number of binding sites differs between the folded and the unfolded states [27]. The binding model introduces two new variables to be fitted compared to the LEM approach, namely the change in denaturant binding sites as the protein unfolds and the binding constants of the denaturant molecules [28]. In a third model, the effect of denaturant on the protein has been described by transfer free energies, however the model is limited in the uncertainties of the free transfer energy for all side chains and the backbone [29,30]. The LEM approach still remains the most applied method for describing the stability of a protein by chaotropes. A linear correlation is observed between the *m*-value and the change in accessible surface area (ASA) as the protein unfolds. Different linear correlations are observed depending on the type of denaturant and whether or not disulphide bonds and crosslinks are present in the native protein, since the presence of crosslinks will result in a more compact denatured state reducing the change in ASA [31]. Because the change in ASA depends on the size of the protein, *m*-values may often be estimated from the length of a protein [32].

The denaturant dependence of the fluorescence signals for the native and the denatured states are often found to be linear [21]:

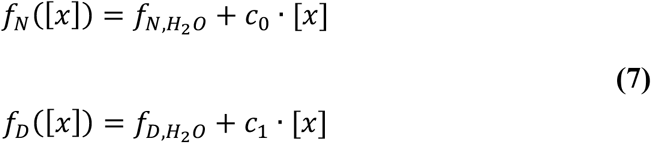

Analogous to the derivation of the expression for the temperature dependence of the spectroscopic signal we get for denaturation by denaturants:

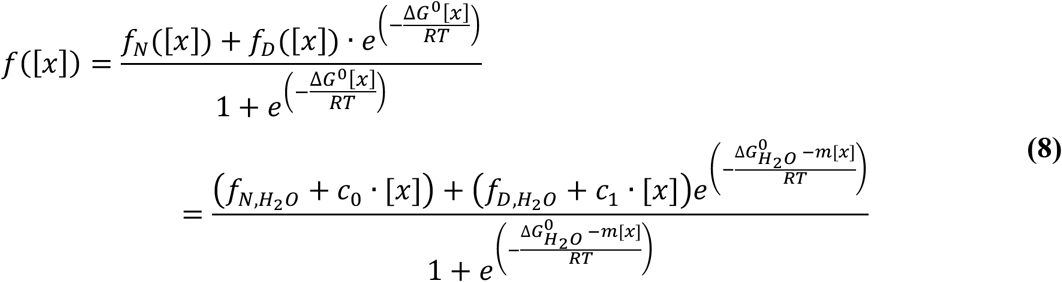

#### Combined temperature and chemical denaturation

To arrive at a combined expression that describes the stability variation as a function of both temperature and the concentration of denaturant we have used that the *m*-value may vary with the temperature as previously observed for other proteins [28,33,34]. We have only included a linear term as we saw no improvement in the fit by including a quadratic term as previously suggested [28].

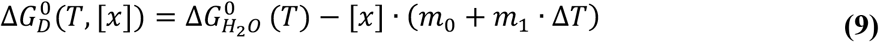

where Δ*T* again is Δ*T* = *T – T_ref_* and *T_ref_* is the temperature where the *m*-value equals *m*_0_. Assuming that *C_p_* is independent of the denaturant concentration equation 5, 8 and 9 can be combined to give an expression that describes the signal:

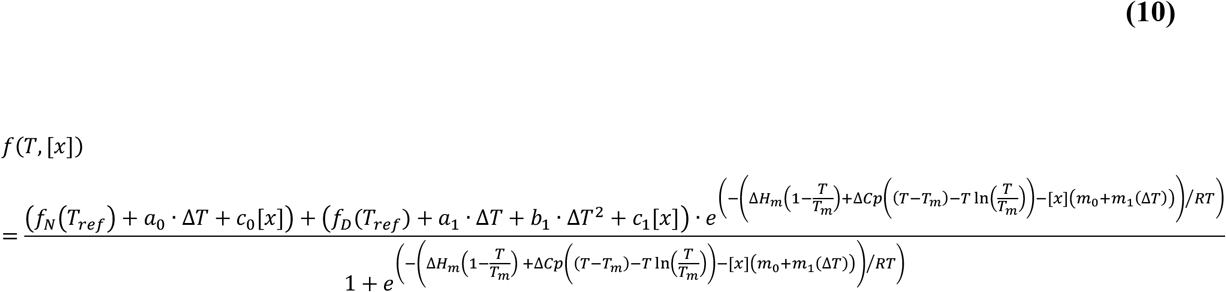

This equation describes the dependency of a spectroscopic signal as a function of the temperature and the denaturant concentration. It contains five thermodynamic parameters (Δ*H*_m_, Δ*C*_p_, *m*_0_, *m*_1_, *T*_m_) that describe the unfolding of the protein, and seven parameters that together describe the temperature and denaturant dependencies of the pre- and post-denaturation baselines (*f*_N_(*T*_ref_), *f*_D_(*T*_ref_), *a*_0_, *a*_1_, *b*_1_, *c*_0_ and *c*_1_).

## Methods

### Protein preparation

Bovine ACBP was transformed in *Escherichia coli* BL21(DE3)/pLysS and grown in LB media supplemented with 100 μg mL^-1^ ampicillin and 50 μg mL^-1^ chloramphenicol. Protein expression was induced by 0.4 mM IPTG at 37 °C for four hours. Cells were harvested by centrifugation (5000xg) for 15 min at 4 °C. The cell paste was resuspended in 20 mM Tris-HCl pH 8, 20 mM DTT and incubated at 30 °C to induce cell lysis by T7 lysozyme. 50 μg mL^-1^ DNase was added and incubated on ice before centrifugation (20,000xg) for 20 min at 4 °C. The pH of the supernatant was adjusted to pH 4 with acetic acid before centrifugation (20,000xg) for 20 min at 4 °C. The pH of the supernatant was adjusted to pH 8 with sodium hydroxide and applied onto a superdex75 26/60 equilibrated with 20 mM Tris-HCl pH 8. The peak with ACBP was collected and applied onto a Q Sepharose HP column equilibrated with 20 mM Tris-HCl pH 8 and eluted with 20 mM Tris-HCl pH 8, 0.5 M NaCl. The peak with ACBP was collected and pH was adjusted to pH 2 with TFA and applied onto a Zorbax C-18 RP-HPLC column equilibrated with 0.1 % TFA and eluted with a linear gradient of 10-80 % acetonitrile in 0.1 % TFA. The eluted ACBP was collected and lyophilized before it was dissolved in 20 mM sodium acetate pH 5.3.

CI2 was transformed into *Escherichia coli* BL21(DE3)/pLysS and grown in LB media supplemented with 100 μg mL^-1^ ampicillin and 50 μg mL^-1^ chloramphenicol. Protein expression was induced by 0.4 mM IPTG at 37 °C for four hours. Cells were harvested by centrifugation (5,000xg) for 15 min at 4 °C. The cell paste was resuspended in 10 mM sodium acetate pH 4.4 followed by centrifugation (20,000xg) for 30 min at 4 °C. The supernatant was diluted with 10 mM sodium acetate pH 4.4 to lower the conductivity before the sample was applied onto a ReSource S column equilibrated with 20 mM sodium acetate pH 4.4. The column was eluted with 20 mM sodium acetate pH 4.4, 1 M NaCl. Fractions with CI2 were collected and dialyzed against water before applied onto a superdex75 16/60 column equilibrated with 20 mM sodium phosphate pH 7.4, 150 mM NaCl. The eluted sample was lyophilized before resuspended in two different buffers, 10 mM glycine-HCl pH 3.0 and 50 mM MES pH 6.25.

### Differential scanning calorimetry

The DSC experiment was performed on a MicroCal VP-DCS at a temperature scan rate of 1 °C/min. Initially, repetitive scans of the background (20 mM Na-acetate, 0.2 M GuHCl, pH 5.3) were performed until the instrument had stabilized (typically 20 hours). The sample was exchanged with 0.04 mg/ml ACBP in 20 mM Na-acetate, 0.2 M GuHCl, pH 5.3 and temperature scans were recorded at 1 °C/min.

### Equilibrium unfolding

Protein concentrations were determined on a Lambda 40 UV/VIS spectrophotometer (PerkinElmer Instrument) using the extinction coefficients of 6,990 M^-1^cm^-1^ for CI2 and 15,470 M^-1^cm^-1^ for ACBP. The protein concentration was adjusted to 10 μM. For denaturant dilution series, two protein stocks were prepared in the appropriate buffer; one without denaturant and another with the maximal concentration of guanidine hydrochloride (GuHCl). Protein samples with different GuHCl concentrations were prepared by mixing different volumes of the two protein stocks to a final volume of 15 μl. A total of 16 samples were prepared with linear increase in [GuHCl]. Fluorescence at 330 nm and 350 nm upon excitation at 280 nm were measured on a Prometheus NT.48 (nanoTemper Technologies) using Prometheus NT.48 high sensitivity capillaries. The temperature range was 15-95 °C with a temperature increment of 1°C /min. For each protein the protein unfolding is determined in triplicates. Global analysis of temperature and solvent denaturation was performed using the lmfit function in python, where the parameters are determined using a least-square curvefitting approach. A Python Jupyter notebook (https://jupyter.org/) with examples for analysing unfolding experiments using the equations and approaches described in this paper is available from https://github.com/KULL-Centre/papers/edit/master/2020/global-analysis-Hamborg-et-al/.

## Results and discussion

To demonstrate the use of the two-dimensional fitting procedure we have measured combined temperature and solvent denaturation of ACBP at pH 5.3 and CI2 at pH 3 and pH 6.25, by following the change in intrinsic Trp fluorescence at 330 nm and 350 nm. For a series of 16 samples of each protein in GuHCl, the temperature was ramped from 15 °C to 95 °C.

The raw fluorescence data are shown in Figure 1. With the Prometheus NT.48 instrument the samples are loaded into cylindrical capillaries and the fluorescence signals from the samples are measured directly on the capillaries. In our experience, the fluorescence intensity from identical samples may vary by up to 10% depending on the position of the capillary in the sample tray. To correct for this, we have normalized the signals in each measurement series to the signal at a temperature where all samples in the series are unfolded. The manipulation corrects for the sample-position-dependent variation. However, it will also remove any putative GuHCl dependence of the fluorescence signal at the wavelength of normalization. This is estimated to less of an error than the variation from the sample position.

**Figure 1.**
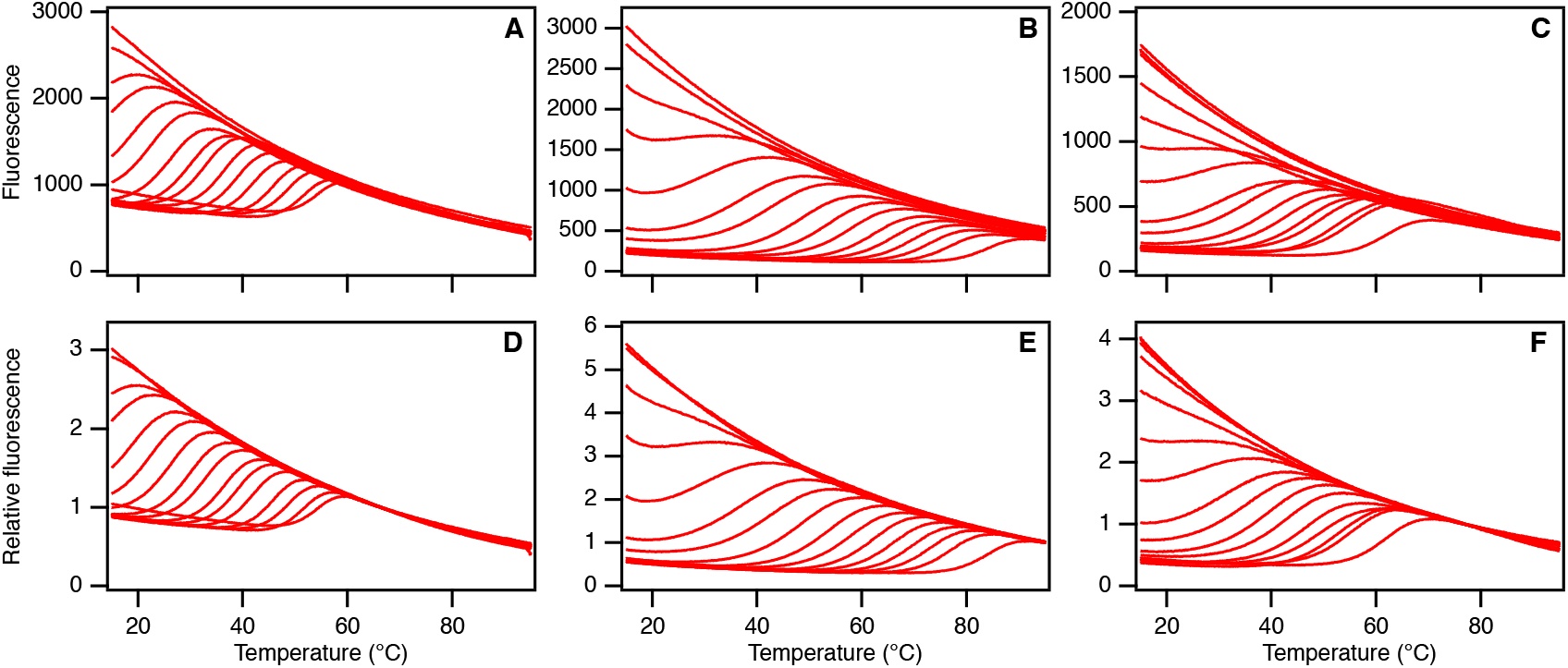
Thermal unfolding curves of ACBP (A+D), CI2 at pH 6.25 (B+E) and CI2 at pH 3.0 (C+F). A-C) Raw fluorescence traces measured at 350 nm with [GuHCl] varying from 0 M to 6 M (curves right to left). D-F) Same traces as in A-C after normalizing to the fluorescence of the unfolded state.

Initially, we fitted the data at each individual temperature to Eq.8. From this analysis we get stability curves with well determined baselines from 20 °C to 35 °C for ACBP, from 30 to 60 °C, for CI2 at pH 6.25 and from 15 °C to 35 °C for CI2 at pH 3.0. The dependence of temperature on Δ*G*^0^, the *m*-value, and *c*_0_ and *c*_1_ (the slopes of the folded and unfolded baselines) are evaluated from these fits (Figure 2 and Figure S1). For all proteins Δ*G*^0^ decreases with temperature as expected. The variation of the *m*-values with temperature is small for both CI2 conditions, whereas the *m*-value for ACBP show a stronger dependence of temperature as previously reported for other proteins [28]. The slopes of the pre- and post-transition baselines, *c*_0_ and *c*_1_, appears to vary little with temperature for ACBP and CI2 at pH 6. For CI2 at pH 3.0, however there appears to be some variation of these parameters with temperature. Two-dimensional fits of the data for CI2 at pH 3.0 to Eq. 10 and to the same equation including temperature variation of *c*_0_ and *c*_1_ give results that are almost identical. Therefore, we have assumed the slopes/curvature in the temperature dimension is independent of GuHCl and *vice versa.* We have modelled the baselines of the folded and the unfolded states as linearly dependent on [GuHCl]. Similarly, the baseline of the fluorescence signal from the folded state is linearly dependent on the temperature, whereas the baseline of the unfolded state has a curved temperature dependence. We have modelled this curvature as a quadratic term, as previously suggested in the literature [22].

**Figure 2.**
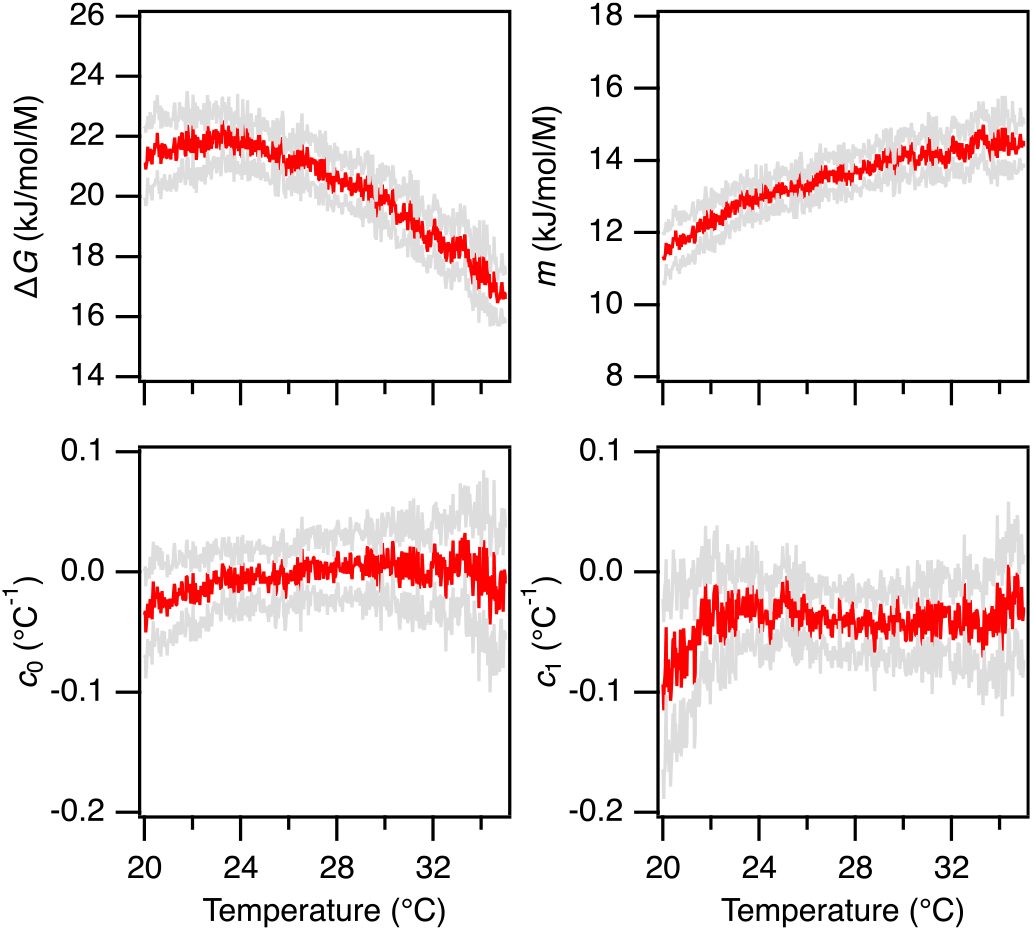
Temperature variation in parameters from fits of fluorescence vs [GuHCl] for ACBP at pH 5.3. Red points represent the values from fits to Eq. 8 and the grey points represent ± standard deviation from the fit.

The fits of the three normalized datasets to Eq. 10 are shown in Figure 3 and summarized in Table 1. The fluorescence of ACBP decreases strongly with addition of salt up to 0.2 M [35] and the baseline at 0 M GuHCl thus does not coincide with baselines at higher concentrations of GuHCl. For CI2 at both pH values we observe that the temperature unfolding is concentration dependent in the absence of GuHCl. Therefore, the data at 0 M GuHCl were excluded for the three denaturation series. For the samples where GuHCl is present the fits reproduce the data well both in the temperature dimension (Figure 3, upper row) and in the GuHCl dimension (Figure 3, lower row). The largest deviations between the data and the fits are observed at low temperatures and high concentrations of GuHCl. We fit the data from the three data series without and with a linear temperature dependence of the *m*-value.

**Figure 3.**
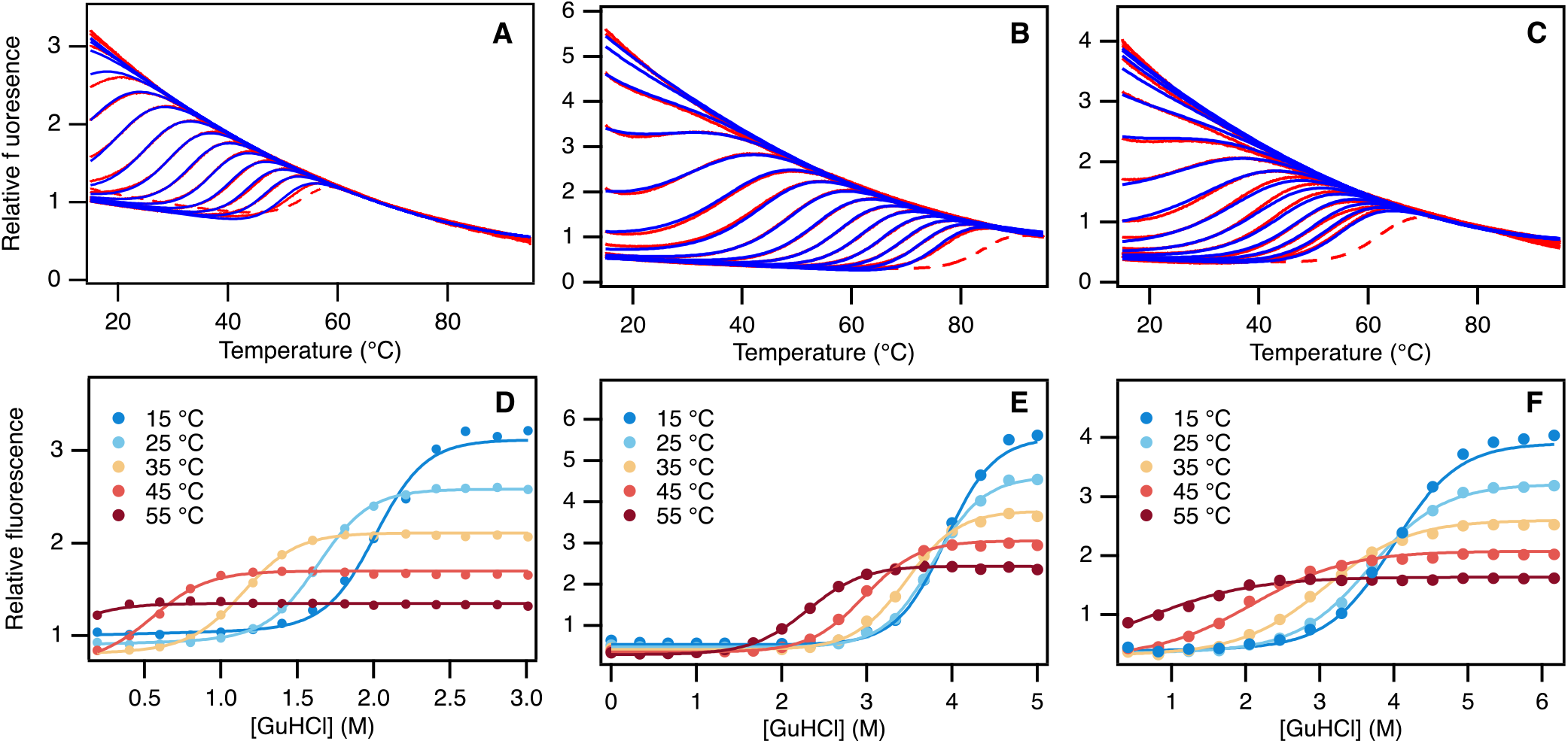
Two-dimensional fits of equilibrium unfolding by temperature and denaturant of ACBP (A+D), CI2 at pH 6.25 (B+E) and CI2 at pH 3.0 (C+F). **A-C)** Temperature unfolding of 15 samples with [GuHCl] varying from 0.3 to 6 M. Fits to Eq. 10 are shown as blue lines on top of the red experimental data. The samples with 0 M GuHCl were not included in the fits and are shown as red dashed lines in panels. **D-F)** GuHCl unfolding curves extracted at five temperatures from panels A-C. The experimental data are shown as filled circles and the fits to Eq. 10 are shown as solid lines.

**Table 1.** Results of two-dimensional fits of protein unfolding curves from fluorescence data. The values are the averages and standard deviations of three independent experiments.

| Sample | $T_m$<br>(K)<sup>a</sup> | $\Delta H_m$<br>(kJ mol<sup>-1</sup>) | $\Delta C_p$<br>(kJ mol<sup>-1</sup> K<sup>-1</sup>) | $m_0$<br>(kJ mol<sup>-1</sup> M<sup>-1</sup>) | $m_1$<br>(kJ mol<sup>-1</sup> M<sup>-1</sup> K<sup>-1</sup>) | $\Delta G_{H_2O}^0$<br>(kJ mol<sup>-1</sup>)<sup>b</sup> |
| --- | --- | --- | --- | --- | --- | --- |
| CI2 pH 3.0<sup>c</sup> | 332.9 ± 0.4 | 249 ± 22 | 4.7 ± 0.5 | 4.6 ± 0.2 |  | 17.2 ± 1.3 |
| CI2 pH 3.0 | 332.5 ± 0.6 | 239 ± 12 | 4.0 ± 0.2 | 4.6 ± 0.4<sup>d</sup> | -0.013 ± 0.013 | 17.2 ± 1.3 |
| CI2 pH 6.25<sup>c</sup> | 352.3 ± 0.2 | 322 ± 11 | 4.3 ± 0.2 | 8.2 ± 0.1 |  | 30.5 ± 0.8 |
| CI2 pH 6.25 | 352.3 ± 0.1 | 322 ± 9 | 4.3 ± 0.3 | 8.2 ± 0.2<sup>d</sup> | 0.0004 ± 0.0045 | 30.5 ± 0.8 |
| ACBP pH 5.3<sup>c</sup> | 326.2 ± 0.1 | 345 ± 15 | 4.6 ± 0.6 | 14.8 ± 0.3 |  | 24.1 ± 0.6 |
| ACBP pH 5.3 | 327.1 ± 0.3 | 390 ± 1 | 8.4 ± 0.8 | 14.6 ± 0.7<sup>d</sup> | 0.13 ± 0.05 | 23.5 ± 0.9 |

<sup>a</sup>In the absence of denaturant.
<sup>b</sup>Value calculated at 298 K.
<sup>c</sup>The linear dependence of the $m$ -value, $m_1$ , is not included in the fit.
<sup>d</sup>Value at $T_{ref} = 298$ K

As seen from Table 1 the fits for CI2 are identical within the error. For ACBP we get an unreasonably high fitted value for Δ*C*_p_ when the *m*-value is allowed to vary with temperature. We thus kept the *m*-value independent of the temperature in the remaining analysis.

To assess the robustness of the fits and the correlations between the fitted parameters, we repeated the fits for each of the three proteins with reduced datasets where we systematically excluded two of the 15 samples from the fit. For each protein we get 15*14/2=105 estimates of the thermodynamically parameters. Inspection of the correlation plots (Figure S2, Figure S3 and Figure S4) shows that the correlation between the parameters vary from protein to protein. *T*m appears to be the parameter least correlated with other parameters whereas Δ*H*^°^ and the *m*-value are most correlated in all datasets. The parameter correlations observed from our analysis are less pronounced than what is seen for direct fits of Δ*G*° and the *m*-value in other work [36]. The average parameters from this analysis are listed in Table S1.

Denaturant unfolding has previously been reported at 298 K for ACBP at pH 5.3 (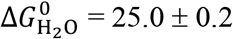 kJ/mol and *m* = 14.7 ± 0.1 kJ/mol/M) [37] and CI2 at pH 6.25 (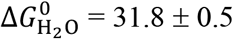 kJ/mol and *m* = 7.9 ± 0.1 kJ/mol/M) [13]. Thermal unfolding has not previously been reported for ACBP. For CI2 DSC experiments have been performed at pH 2.8 and pH 3.2 on a variant of CI2 including a disordered N-terminal tail of 19 amino acid residues [6]. In the absence of denaturant, *T*_m_ is 328 K and 337 K at pH 2.8 and 3.2, respectively. This is in good agreement with *T*_m_ = 333 K that we find at pH 3, despite that the sample without GuHCl was not included in our fit *(vide supra). ΔH* = 249 kJ/mol found in this work at pH 3 is also in between the values of 221 kJ/mol and 255 kJ/mol previously reported at pH 2.8 and 3.2, respectively. In contrast, Δ*C*_p_ = 4.7 kJ/mol found here is more than 40% different from the value of 3.3 kJ/mol previously reported. In summary, *T*_m_, 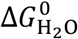, and *m* determined by our method are in good agreement with previous values. Δ*H* and Δ*C*_p_ cannot directly be compared to the published values, which were recorded on CI2 including a tail of 19 disordered amino acid residues. We attempted to perform DSC experiments on both CI2 and ACBP to complement our data. CI2 precipitates at high temperatures and ACBP does not behave as two-state at high temperatures and at a protein concentration of 0.2 mg/ml that is needed for proper signal to noise in the DSC experiment. At 0.04 mg/ml and in the presence of 0.2 M GuHCl, DSC was measured on ACBP. The baseline of the folded state, however, is not well determined impairing analysis of the data (Figure S5).

The additional information acquired by adding a temperature dimension to denaturant unfolding should constrain the data fitting better than an equilibrium folding curve recorded at only a single temperature, and in addition also provide estimates of *T*_m_, Δ*H*_m_ and Δ*C*_p_. We therefore tested how few data points in the GuHCl dimension were needed to still get reliable results from the analysis. Based on the 15 samples we included in the fits above for each protein, we generated subsets of data by selecting every second, every third and every fourth sample resulting in datasets with 8, 5 and 4 samples, respectively evenly spaced in the GuHCl concentration range. The results of fitting these datasets to Eq. 10 are shown in Table 2. In general, the results are very similar demonstrating that the fits are well constrained by samples at only a few GuHCl concentrations. In principle, except for the *m*-value, a single thermal unfolding curve should be enough to fit *T*_m_, Δ*H*_m_ and Δ*C*_p_. In reality, we find that the baselines for the folded and unfolded states are not well constrained from a single sample and that four or five samples at well separated denaturant concentrations are needed. However, care should be taken if using only a few samples as a single sample that was not accurately prepared with precise denaturant and protein concentrations will have a large influence on the fitted parameters.

**Table 2.** Thermodynamic parameters from fits of reduced datasets. The listed values are the averages and standard deviations of three independent experiments.

| | No of samples | $T_m$<br>(K)<sup>a</sup> | $\Delta H_m$<br>(kJ mol<sup>-1</sup>) | $\Delta C_p$<br>(kJ mol<sup>-1</sup> K<sup>-1</sup>) | m<br>(kJ mol<sup>-1</sup> M<sup>-1</sup>) | $\Delta G_{H_2O}^0$<br>(kJ mol<sup>-1</sup>)<sup>b</sup> |
| --- | --- | --- | --- | --- | --- | --- |
| CI2, pH 3 | 15 points | 332.9 ± 0.4 | 249 ± 22 | 4.7 ± 0.5 | 4.6 ± 0.2 | 17.1 ± 1.2 |
|  | 8 points | 332.8 ± 0.5 | 244 ± 8 | 4.6 ± 0.4 | 4.5 ± 3 | 16.8 ± 0.4 |
|  | 5 points | 332.5 ± 0.6 | 237 ± 3 | 4.5 ± 0.3 | 4.4 ± 0.2 | 16.3 ± 0.4 |
|  | 4 points | 332.3 ± 0.5 | 264 ± 13 | 5.1 ± 0.2 | 4.7 ± 0.1 | 17.9 ± 1.2 |
| CI2, pH 6 | 15 points | 352.3 ± 0.2 | 322 ± 11 | 4.3 ± 0.2 | 8.2 ± 0.1 | 30.6 ± 0.8 |
|  | 8 points | 352.3 ± 0.1 | 328 ± 3 | 4.4 ± 0.1 | 8.4 ± 0.2 | 31.0 ± 0.1 |
|  | 5 points | 352.5 ± 0.2 | 332 ± 33 | 4.5 ± 0.6 | 8.5 ± 0.6 | 31 ± 3 |
|  | 4 points | 352.4 ± 0.5 | 340 ± 10 | 4.7 ± 0.2 | 8.6 ± 0.3 | 31.7 ± 0.8 |
| ACBP | 15 points | 326.2 ± 0.1 | 345 ± 15 | 4.6 ± 0.6 | 14.8 ± 0.3 | 24.1 ± 0.6 |
|  | 8 points | 326.3 ± 0.1 | 341 ± 14 | 4.5 ± 0.5 | 14.7 ± 0.3 | 24.0 ± 0.6 |
|  | 5 points | 326.5 ± 0.1 | 346 ± 23 | 4.5 ± 0.8 | 14.8 ± 0.6 | 24.4 ± 0.6 |
|  | 4 points | 326.4 ± 0.1 | 358 ± 4 | 5.0 ± 0.4 | 15.2 ± 0.4 | 24.9 ± 0.5 |

<sup>a</sup>In the absence of denaturant.
<sup>b</sup>Calculated using Eq. 2 at 298 K

## Conclusion

In this work we have demonstrated that well-determined values 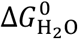 for ACBP and CI2 can be extracted from as little as four temperature-unfolding curves at different GuHCl concentrations. Combining this with the low sample requirements in the nanoDSF instrument, allows high quality stability data to be acquired with less than 10 μg protein. Such amounts can often be purified in 96 well plate formats allowing high throughput analysis of proteins from mutant libraries and to identify protein variants with change in stability. For ACBP and CI2 thermodynamic parameters for the folding process cannot be determined by DSC and the method we present here provides an alternative for these and other proteins that do not behave well in DSC. Python code to visualize and fit data to the models we describe is available online, and can easily be modified to include other parameterizations of the baselines or thermodynamic models.

## Abbreviations

ACBP: bovine acyl-coenzyme A binding protein;
ASA: accessible surface area;
CI2: barley chymotrypsin inhibitor 2;
DSC: differential scanning calorimetry;
LEM: linear extrapolation model.

## Acknowledgements

This work was supported by the Novo Nordisk Foundation [grant number NNF150C0016360]. The authors thank Pia Skovgaard for technical assistance. KLL and KT are members of Integrative Structural Biology at the University of Copenhagen (www.isbuc.ku.dk).

## References

[1] C.M. Johnson, Differential scanning calorimetry as a tool for protein folding and stability, Arch. Biochem. Biophys. 531 (2013) 100–109. https://doi.org/10.1016/j.abb.2012.09.008.

[2] N.J. Greenfield, Using circular dichroism collected as a function of temperature to determine the thermodynamics of protein unfolding and binding interactions, Nat. Protoc. 1 (2007) 2527–2535. https://doi.org/10.1038/nprot.2006.204.

[3] F.H. Niesen, H. Berglund, M. Vedadi, The use of differential scanning fluorimetry to detect ligand interactions that promote protein stability, Nat. Protoc. 2 (2007) 2212–2221. https://doi.org/10.1038/nprot.2007.321.

[4] M.R. Eftink, Fluorescence techniques for studying protein structure., Methods Biochem. Anal. 35 (1991) 127–205. https://doi.org/10.1002/9780470110560.ch3.

[5] D.E. Otzen, L.S. Itzhaki, N.F. ElMasry, S.E. Jackson, A.R. Fersht, Structure of the transition state for the folding/unfolding of the barley chymotrypsin inhibitor 2 and its implications for mechanisms of protein folding., Proc. Natl. Acad. Sci. U. S. A. 91 (1994) 10422–5. https://doi.org/10.1073/pnas.91.22.10422.

[6] S.E. Jackson, A.R. Fersht, Folding of chymotrypsin inhibitor 2. 1. Evidence for a two-state transition., Biochemistry. 30 (1991) 10428–35. https://doi.org/10.1021/bi00107a010.

[7] S.E. Jackson, A.R. Fersht, Contribution of Residues in the Reactive Site Loop of Chymotrypsin Inhibitor 2 to Protein Stability and Activity, Biochemistry. 33 (1994) 13880–13887. https://doi.org/10.1021/bi00250a042.

[8] D.E. Otzen, M. Rheinnecker, A.R. Fersht, Structural Factors Contributing to the Hydrophobic Effect: The Partly Exposed Hydrophobic Minicore in Chymotrypsin Inhibitor, Biochemistry. 34 (1995) 13051–13058. https://doi.org/10.1021/bi00040a016.

[9] S.E. Jackson, M. Moracci, N. ElMasry, C.M. Johnson, A.R. Fersht, Effect of cavity-creating mutations in the hydrophobic core of chymotrypsin inhibitor 2., Biochemistry. 32 (1993) 11259–69. https://doi.org/10.1021/bi00093a001.

[10] A.G. Ladurner, L.S. Itzhaki, A.R. Fersht, Strain in the folding nucleus of chymotrypsin inhibitor 2, Fold. Des. 2 (1997) 363–368. https://doi.org/10.1016/S1359-0278(97)00050-3.

[11] C. Lawrence, J. Kuge, K. Ahmad, K.W. Plaxco, Investigation of an Anomalously Accelerating Substitution in the Folding of a Prototypical Two-State Protein, J. Mol. Biol. 403 (2010) 446–458. https://doi.org/10.1016/jjmb.2010.08.049.

[12] Y.P. Pan, V. Daggett, Direct comparison of experimental and calculated folding free energies for hydrophobic deletion mutants of chymotrypsin inhibitor 2: Free energy perturbation calculations using transition and denatured states from molecular dynamics simulations of unfoldi, Biochemistry. 40 (2001) 2723–2731. https://doi.org/10.1021/bi0022036.

[13] L.S. Itzhaki, D.E. Otzen, A.R. Fersht, The structure of the transition state for folding of chymotrypsin inhibitor 2 analysed by protein engineering methods: evidence for a nucleation-condensation mechanism for protein folding., J. Mol. Biol. 254 (1995) 260–88. https://doi.org/10.1006/jmbi.1995.0616.

[14] J.K. Thomsen, B.B. Kragelund, K. Teilum, J. Knudsen, F.M. Poulsen, Transient intermediary states with high and low folding probabilities in the apparent two-state folding equilibrium of ACBP at low pH., J. Mol. Biol. 318 (2002) 805–14. https://doi.org/10.1016/S0022-2836(02)00159-6.

[15] B.B. Kragelund, P. Osmark, T.B. Neergaard, J. Schiødt, K. Kristiansen, J. Knudsen, F.M. Poulsen, The formation of a native-like structure containing eight conserved hydrophobic residues is rate limiting in two-state protein folding of ACBP., Nat. Struct. Biol. 6 (1999) 594–601. https://doi.org/10.1038/9384.

[16] B.B. Kragelund, P. Højrup, M.S. Jensen, C.K. Schjerling, E. Juul, J. Knudsen, F.M. Poulsen, Fast and one-step folding of closely and distantly related homologous proteins of a four-helix bundle family., J. Mol. Biol. 256 (1996) 187–200. https://doi.org/10.1006/jmbi.1996.0076.

[17] K. Teilum, T. Thormann, N.R. Caterer, H.I. Poulsen, P.H. Jensen, J. Knudsen, B.B. Kragelund, F.M. Poulsen, Different secondary structure elements as scaffolds for protein folding transition states of two homologous four-helix bundles, Proteins Struct. Funct. Bioinforma. 59 (2005) 80–90. https://doi.org/10.1002/prot.20340.

[18] K.L. Maxwell, D. Wildes, A. Zarrine-Afsar, M.A. De Los Rios, A.G. Brown, C.T. Friel, L. Hedberg, J.-C. Horng, D. Bona, E.J. Miller, A. Vallée-Bélisle, E.R.G. Main, F. Bemporad, L. Qiu, K. Teilum, N.-D. Vu, A.M. Edwards, I. Ruczinski, F.M. Poulsen, B.B. Kragelund, S.W. Michnick, F. Chiti, Y. Bai, S.J. Hagen, L. Serrano, M. Oliveberg, D.P. Raleigh, P. Wittung-Stafshede, S.E. Radford, S.E. Jackson, T.R. Sosnick, S. Marqusee, A.R. Davidson, K.W. Plaxco, Protein folding: Defining a “standard” set of experimental conditions and a preliminary kinetic data set of two-state proteins, Protein Sci. 14 (2005) 602–616. https://doi.org/10.1110/ps.041205405.

[19] Q. Yi, M.L. Scalley, K.T. Simons, S.T. Gladwin, D. Baker, Characterization of the free energy spectrum of peptostreptococcal protein L, Fold. Des. 2 (1997) 271–280. https://doi.org/10.1016/S1359-0278(97)00038-2.

[20] W.J. Becktel, J.A. Schellman, Protein stability curves, Biopolymers. 26 (1987) 1859–1877. https://doi.org/10.1002/bip.360261104.

[21] M.R. Eftink, The use of fluorescence methods to monitor unfolding transitions in proteins, Biophys. J. 66 (1994) 482–501. https://doi.org/10.1016/S0006-3495(94)80799-4.

[22] K. Saini, S. Deep, Relationship between the wavelength maximum of a protein and the temperature dependence of its intrinsic tryptophan fluorescence intensity, Eur. Biophys. J. 39 (2010) 1445–1451. https://doi.org/10.1007/s00249-010-0601-3.

[23] K. Lindorff-Larsen, J.R. Winther, Surprisingly high stability of barley lipid transfer protein, LTP1, towards denaturant, heat and proteases, FEBS Lett. 488 (2001) 145–148. https://doi.org/10.1016/S0014-5793(00)02424-8.

[24] M.M. Santoro, D.W. Bolen, Unfolding free energy changes determined by the linear extrapolation method. 1. Unfolding of phenylmethanesulfonyl.alpha.-chymotrypsin using different denaturants, Biochemistry. 27 (1988) 8063–8068. https://doi.org/10.1021/bi00421a014.

[25] C. Tanford, Protein denaturation. C. Theoretical models for the mechanism of denaturation., Adv. Protein Chem. 24 (1970) 1–95. https://doi.org/10.1016/S0065-3233(08)60241-7.

[26] R.F. Greene, C.N. Pace, Urea and guanidine hydrochloride denaturation of ribonuclease, lysozyme, alpha-chymotrypsin, and beta-lactoglobulin., J. Biol. Chem. 249 (1974) 5388–93. http://www.ncbi.nlm.nih.gov/pubmed/4416801.

[27] J.F. Brandts, The Thermodynamics of Protein Denaturation. II. A Model of Reversible Denaturation and Interpretations Regarding the Stability of Chymotrypsinogen, J. Am. Chem. Soc. 86 (1964) 4302–4314. https://doi.org/10.1021/ja01074a014.

[28] M.E. Zweifel, D. Barrick, Relationships between the temperature dependence of solvent denaturation and the denaturant dependence of protein stability curves., Biophys. Chem. 101-102 (2002) 221–37. https://doi.org/10.1016/s0301-4622(02)00181-3.

[29] C. Tanford, Isothermal Unfolding of Globular Proteins in Aqueous Urea Solutions, J. Am. Chem. Soc. 86 (1964) 2050–2059. https://doi.org/10.1021/ja01064a028.

[30] J.M. Scholtz, G.R. Grimsley, C.N. Pace, Solvent denaturation of proteins and interpretations of the m value., Methods Enzymol. 466 (2009) 549–565. https://doi.org/10.1016/s0076-6879(09)66023-7.

[31] J.K. Myers, C. Nick Pace, J. Martin Scholtz, Denaturant m values and heat capacity changes: Relation to changes in accessible surface areas of protein unfolding, Protein Sci. 4 (1995) 2138–2148. https://doi.org/10.1002/pro.5560041020.

[32] C.D. Geierhaas, A.A. Nickson, K. Lindorff-Larsen, J. Clarke, M. Vendruscolo, BPPred: A Web-based computational tool for predicting biophysical parameters of proteins, Protein Sci. 16 (2006) 125–134. https://doi.org/10.1110/ps.062383807.

[33] D.J. Felitsky, M.T. Record, Thermal and urea-induced unfolding of the marginally stable lac repressor DNA-binding domain: A model system for analysis of solute effects on protein processes, Biochemistry. 42 (2003) 2202–2217. https://doi.org/10.1021/bi0270992.

[34] A. Amsdr, N.D. Noudeh, L. Liu, T. V. Chalikian, On urea and temperature dependences of m-values, J. Chem. Phys. 150 (2019) 215103. https://doi.org/10.1063/1.5097936.

[35] K. Teilum, F.M. Poulsen, M. Akke, The inverted chevron plot measured by NMR relaxation reveals a native-like unfolding intermediate in acyl-CoA binding protein, Proc. Natl. Acad. Sci. U. S. A. 103 (2006) 6877–6882. https://doi.org/10.1073/pnas.0509100103.

[36] K. Lindorff-Larsen, Dissecting the statistical properties of the Linear Extrapolation Method of determining protein stability, (2020) 1–12. http://arxiv.org/abs/2002.01018.

[37] K. Teilum, B.B. Kragelund, F.M. Poulsen, Transient structure formation in unfolded acyl-coenzyme A-binding protein observed by site-directed spin labelling., J. Mol. Biol. 324 (2002) 349–57. https://doi.org/10.1016/s0022-2836(02)01039-2.

