## Supplementary material for "Global analysis of protein stability by temperature and chemical denaturation": Figure S1

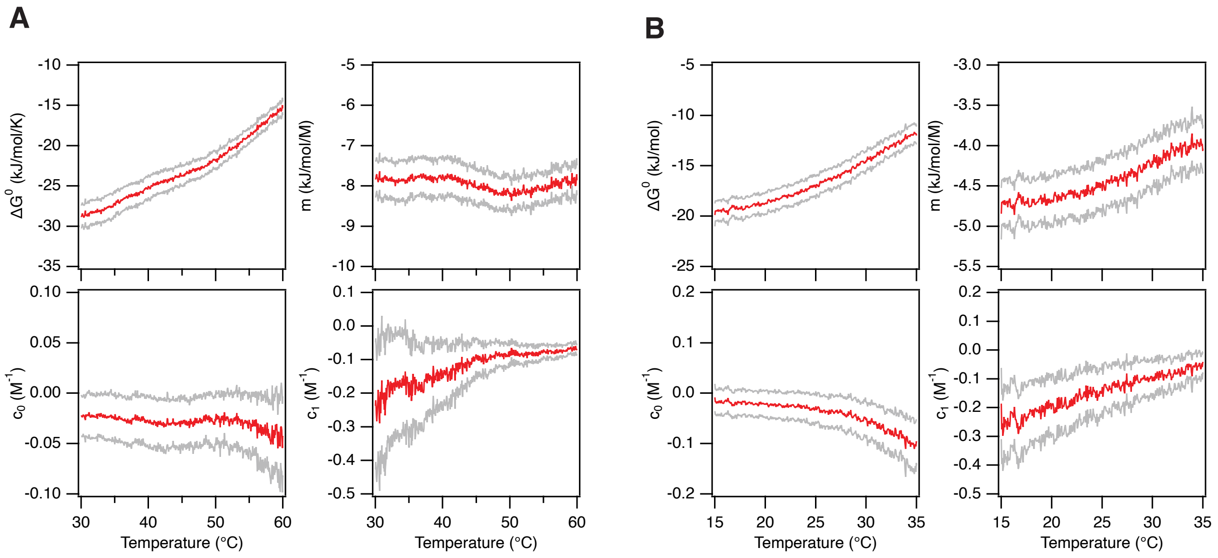


**Figure S1** Temperature variation in parameters from fits of fluorescence vs [GuHCl] for CI2 at (A) pH 6.25 and (B) pH 3.0. Red points represent the values from fits to Eq. 8 and the grey points represent ± standard deviation from the fit.
