## Supplementary figures and images for "Global analysis of protein stability by temperature and chemical denaturation"

### Figure S2

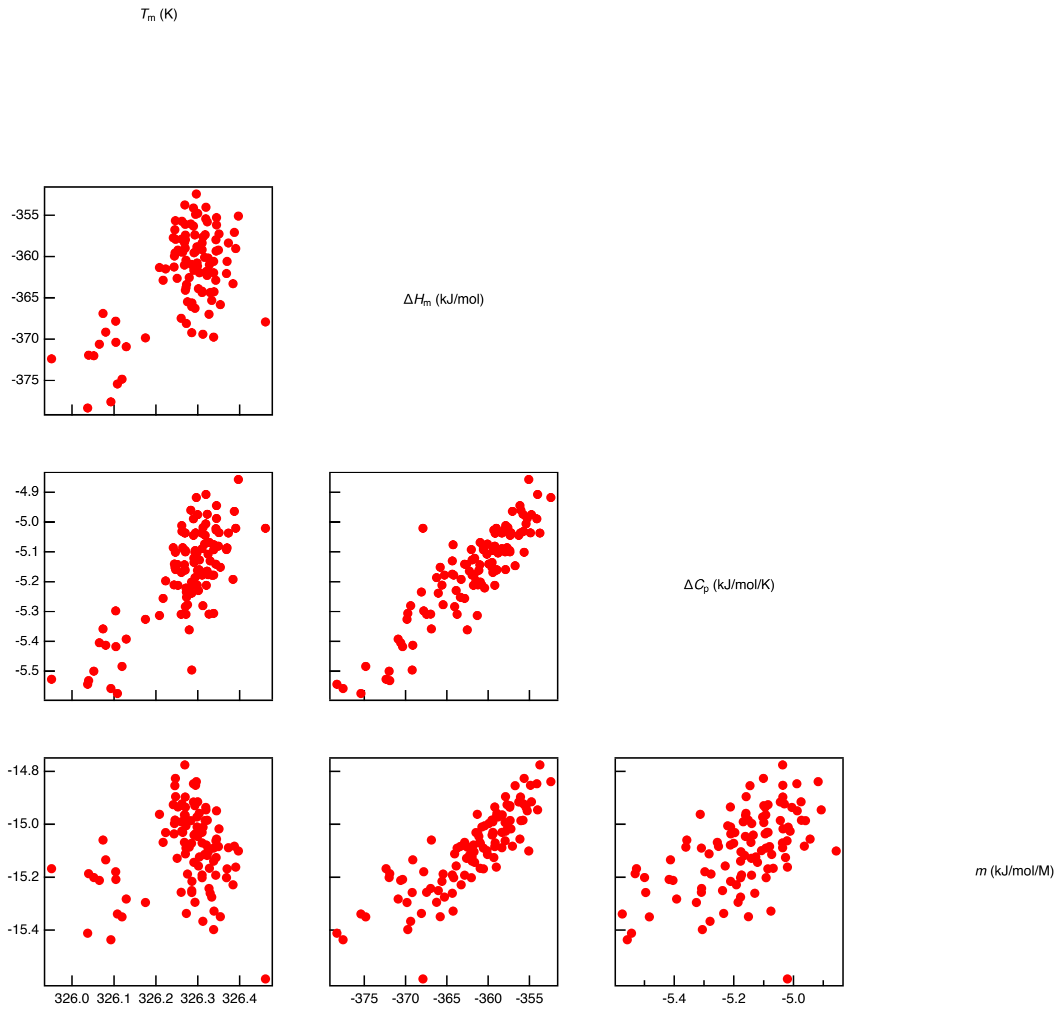


**Figure S2** Correlations between fit parameters from reduced data set of ACBP.

### Figure S3

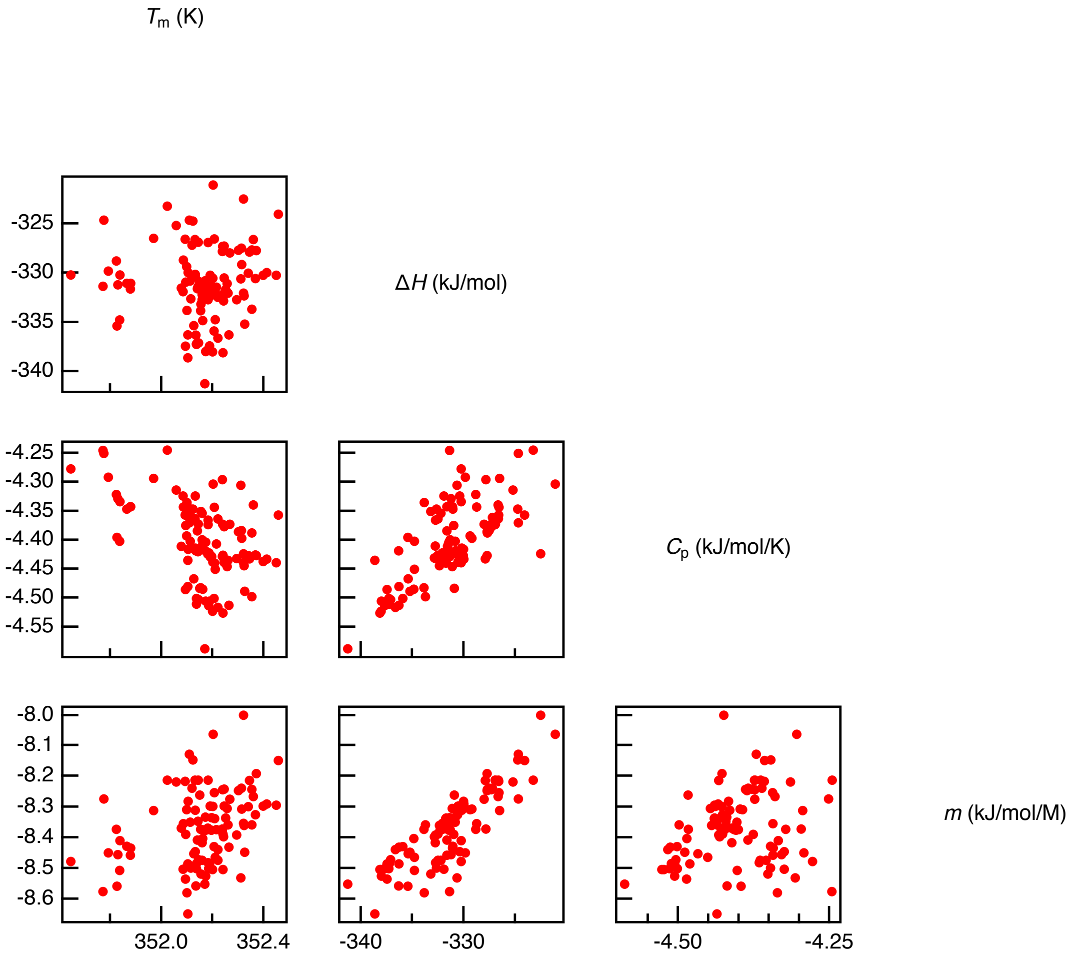


**Figure S3** Correlations between fit parameters from reduced data set of CI2 at pH 6.25.

### Figure S4

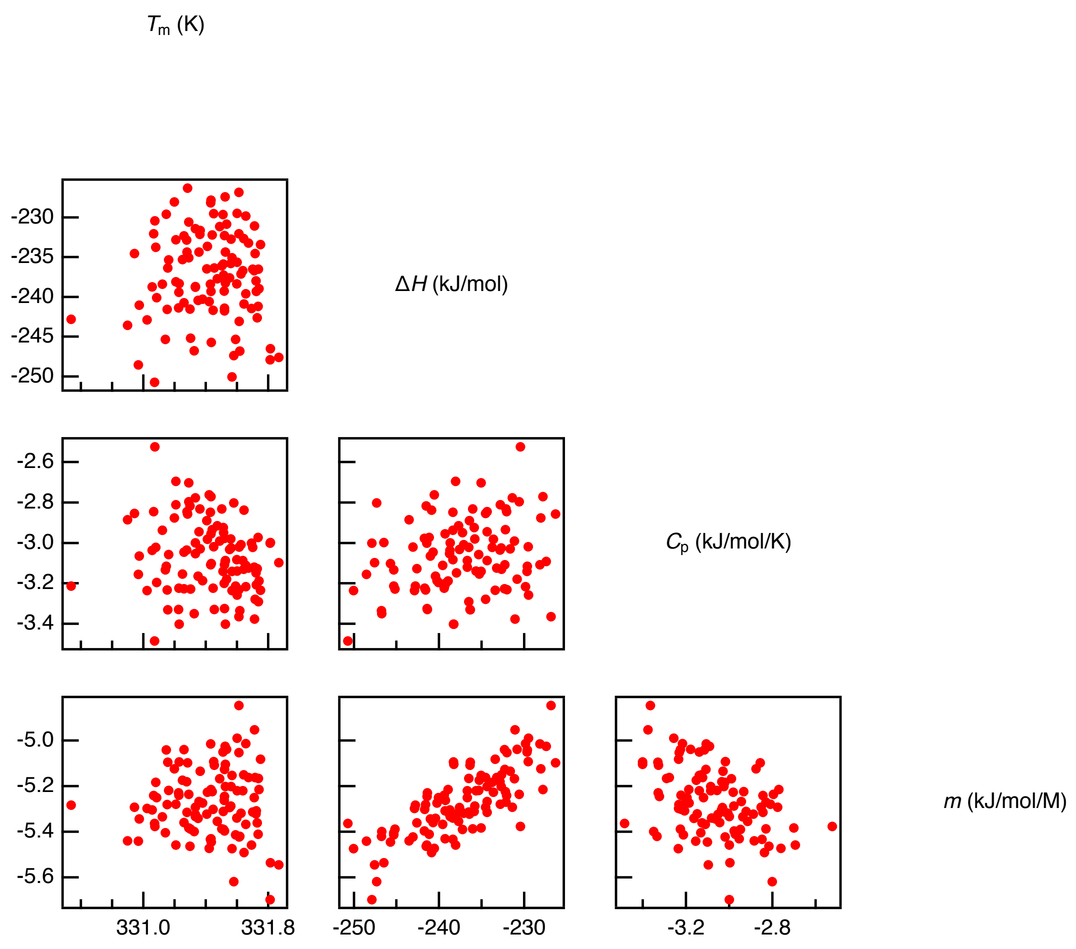


**Figure S4** Correlations between fit parameters from reduced data set of CI2 at pH 3.0.
