## Supplementary material for "Global analysis of protein stability by temperature and chemical denaturation": Figure S5

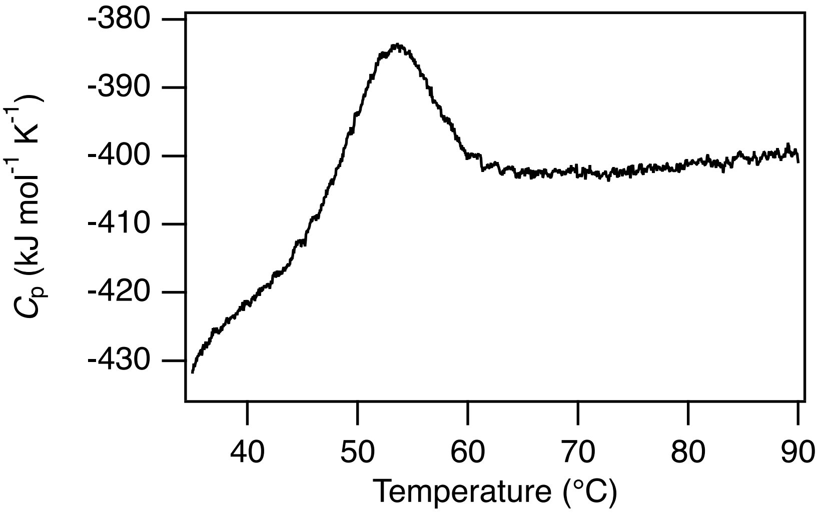


**Figure S5** Differential Scanning Calorimetry of ACBP in 20 mM Na-acetate, 0.2 M GuHCl, pH 5.3. The data have been truncated at 35 °C as the baseline of the folded state is highly curved.
