## Supplementary material for "Global analysis of protein stability by temperature and chemical denaturation": Table S1

**Table S1** Bootstrap analysis of the two-dimensional fit. The robustness of the fitting routine was assessed by fits of 105 reduced datasets in which 2 of out of 15 samples were left out. The average and standard deviation of the fitted parameters were calculated. The analysis was performed for one unfolding experiment on each protein.

| | $T_m$ (K)[a] | $\Delta H_m$ (kJ mol$^{-1}$) | $\Delta C_p$ (kJ mol$^{-1}$ K$^{-1}$) | $m$ (kJ mol$^{-1}$ M$^{-1}$) | $\Delta G^0_{H_2O}$ (kJ mol$^{-1}$)[b] |
|---|---|---|---|---|---|
| CI2, pH 3 | 331.4 ± 0.2 | 237 ± 6 | 3.1 ± 0.2 | 5.3 ± 0.2 | 18.5 ± 0.6 |
| CI2, pH 6 | 352.2 ± 0.2 | 331 ± 4 | 4.4 ± 0.1 | 8.4 ± 0.1 | 30.6 ± 0.8 |
| ACBP | 326.3 ± 0.1 | 362 ± 6 | 5.2 ± 0.2 | 15.1 ± 0.2 | 24.8 ± 0.3 |
